# Downstream Effects: Impact of Antibiotic Pollution on an Aquatic Host-Parasite Interaction

**DOI:** 10.1101/2020.11.07.372813

**Authors:** Hannah G. Melchiorre, Stephanie O. Gutierrez, Dennis J. Minchella, J. Trevor Vannatta

## Abstract

The global increase in antibiotic use has led to contamination of freshwater environments occupied by parasites and their hosts. Despite the identified impacts of antibiotics on humans and wildlife, the effect of antibiotics on host-parasite life cycles is relatively unexplored. We utilize the trematode parasite *Schistosoma mansoni*, and its snail intermediate host *Biomphalaria glabrata* to investigate the influence of an ecologically relevant antibiotic concentration on the life history characteristics of both parasite and host. Our results demonstrate that antibiotics not only accelerate parasite development time, but also increase host reproduction and delay parasite-induced host castration. We propose that antibiotic exposure alters host microbiome composition, leading to increased host susceptibility and higher parasite production. Using a mathematical model, we suggest that life history alterations associated with antibiotics are likely to increase parasite transmission and disease burden. Our study suggests that antibiotic pollution could impact freshwater ecosystems by influencing host-parasite dynamics and potentially increase the burden of schistosomiasis in endemic regions.

## Introduction

Antibiotic usage is increasing worldwide in association with growing demand in livestock production, industry, and human healthcare (Daghrir and Droghui, 2013). As a result, antibiotic contamination from wastewater treatment plants and sewers is often deposited in freshwaters (Kraemer et al., 2019). Presence of antibiotics in aquatic environments affects the life forms within them, extending beyond microscopic organisms to other non-target species (Danner et al., 2019, Sundberg and Karvonen, 2018). Although antibiotic concentrations in freshwater are not typically lethal to non-target organisms, the sublethal impact of runoff on biotic interactions is largely unknown (Cairns et al., 2018, Kim et al., 2014).

The magnitude and severity of antibiotic contamination can have diverse effects depending on the nature of the biotic system (Sundberg and Karvonen, 2018). Antibiotics can act as environmental stressors to alter bacterial communities through direct or indirect mechanisms (Grenni et al., 2018). Antibiotics are designed to decrease pathogenic bacteria within an organism; however, they also impact overall bacterial community diversity within the microbiome and alter community function (Yoon and Yoon, 2018, Morley, 2010, Akbar et al., 2020). Research has shown that the microbiome plays a third-party role in host-parasite interactions (Abraham et al., 2017, Gall et al., 2016, Knutie et al., 2017a, Knutie et al., 2017b), however, few studies have explored the effects of sublethal antibiotic exposure on host-parasite interactions (Morley, 2009).

Antibiotics likely have widespread impacts on many host-parasite systems either directly (by influencing the organisms involved) or indirectly (by influencing predators/prey/parasites of the organisms involved; Pravdová et al., 2020). For instance, decreased immune responses have been reported in pond snails following antibiotic exposure (Gust et al., 2013). Within hosts, reductions in gut microbiome diversity are associated with an increased susceptibility to *S. mansoni* colonization in mice, suggesting a third-party role of microbiome diversity in immune system regulation (Viera et al., 1987). Additionally, immune priming is naturally induced by the gut microbiome during *Plasmodium* infections in *Anopheles gambiae*, and removal of this priming effect resulted in a higher incidence of infection severity and re-infections (Rodrigues et al., 2010). In other systems, an inverse relationship between host, gut microbiota composition and susceptibility to parasite infection has been described, with differences observed in the microbiomes of *S. mansoni*-resistant and susceptible *Biomphalaria* snails (Cortés et al., 2020, Portet et al., 2021). However, the influence of antibiotics in host-parasite systems is not limited to host microbiomes. The effect of antibiotic contamination on parasite microbiomes has also recently been shown (Jorge et al., 2021). Parasite bacterial composition can be indirectly altered by exposing hosts to various antibiotics resulting in divergent influences depending on the antibiotic used (Jorge et al., 2021). Thus, antibiotic effects such as enhanced susceptibility discussed above may be related to altered host or parasite microbiome diversity and/or composition (Pravdová et al., 2020). Understanding the consequences of these altered microbiomes is urgent as contamination of freshwater ecosystems with various pharmaceuticals, such as tetracycline, continues.

Tetracycline is a common broad-spectrum antibiotic that contaminates freshwater environments due to its water solubility and widespread use in agriculture (Lin et al., 2013). The ubiquity of contaminant sources has recently raised concern over increased incidence of antibiotic resistance (Andrade et al., 2020). As such, antibiotic resistance may be used as a proxy to locate overlaps between antibiotic contamination and disease prevalence (Andrade et al., 2020). Despite connections between high rates of tetracycline resistance and prevalence of parasitic disease, little is known about how these interactions impact host-parasite dynamics, such as those seen in schistosomiasis (Faleye et al., 2018).

*Schistosoma mansoni* is a parasitic blood-fluke that causes the human disease schistosomiasis in tropical regions and accounts for as many as 200,000 annual deaths (World Health Organization, 2020). Eggs of *S. mansoni* from infected humans hatch into miracidia, a larval stage of the parasite, when they contact freshwater. These miracidia can penetrate the snail intermediate host, *Biomphalaria glabrata.* Maturation of the parasite occurs within the snail gonads leading to castration of the host. The snails release free-swimming larval stages called cercariae which directly infect humans as the definitive host (CDC, 2018).

Given the human toll of schistosomiasis and the use of antibiotics in medicine and agriculture in tropical regions (Faleye et al., 2018), we designed an experiment to analyze the effects of tetracycline antibiotic contamination on the life history of *S. mansoni* and its snail host. The antibiotic chosen for this experiment was tetracycline due to its water solubility, easy accessibility, and widespread use in agriculture and human medicine (Daghrir and Droghui, 2013). The impact of tetracycline antibiotic exposure on snail growth and reproduction is debated. Some studies show inhibitory effects (Chernin, 1957, Chernin and Schork, 1960), while more recent findings show an antibiotic-induced increase in growth and reproduction (Flaherty and Dodson, 2005, Gaskins et al., 2002). Previous work which demonstrated growth inhibition used dramatically higher antibiotic doses. Our study focuses on the impact of a low dose, ecologically relevant concentration of antibiotics on both host and the parasite. We chose a common concentration found near wastewater treatment plants as residual antibiotic concentrations can fluctuate from 2ng/L to more than 50μg/L depending on location (Islam and Gilbride, 2019, Xu et al., 2021). We predicted that tetracycline would accelerate the development of the parasite, increase parasite reproductive output, and enhance host reproduction potentially via alterations to the host microbiome composition. Our predictions consider a dynamic relationship between immune function and infection susceptibility (Hernández-Gómez et al., 2020, Knutie et al., 2017). Antibiotics influence immune function by modifying bacterial composition within microbiomes (Akbar et al., 2020), which can impact host defense to parasites. We then use our results to parameterize an epidemiological model and demonstrate potential long-term consequences of freshwater antibiotic contamination on human disease burden. Although mechanisms underpinning enhanced host and parasite production due to antibiotic exposure have yet to be fully explored, our results suggest that antibiotic contamination may play a significant role in this host-parasite system.

## Methods

One hundred sixty lab-reared *B. glabrata* snails were used in a full factorial experiment combining parasitic infection and antibiotic exposure for a total of four treatments (antibiotic + parasite, parasite only, antibiotic only, control; Table S1). Forty snails per treatment were size-matched ranging from 8-13mm in shell diameter and housed individually in 120ml jars. Prior to parasite exposure, snails underwent a 4-day acclimation period in well water or well water with the set concentration of antibiotic. Ecologically relevant concentrations of tetracycline vary from region to region, but 50μg/L is a common concentration in areas near waste treatment plants and was used for this experiment (Daghrir and Droghui, 2013, Islam and Gilbride, 2019, Xu et al., 2021). A 50μg/L solution of antibiotic was replaced every 7 days to maintain efficacy and minimize the impact of antibiotic degradation (Schmidt et al., 2007). Antibiotic solutions were prepared by dissolving 1 mg of tetracycline (Research Products International, Tetracycline HCl, Lot #36063-101361) in 20 L of well-water a few hours prior to use.

Each snail in the parasite only treatment and antibiotic + parasite treatment was exposed to 8 miracidia of *S. mansoni* for 24 hours. Unexposed snails were sham exposed for the same period of time. Miracidia were harvested from livers of infected mice (in accordance with Purdue Animal Care and Use Committee protocol #1111000225) by blending in saline and filtering according to standard protocols (Tucker et al., 2013). To quantify host reproductive output, egg masses were counted and removed from all individual snail housing jars weekly for 9 weeks. *Biomphalaria* snails are hermaphroditic and can self-fertilize but may also store sperm from previous encounters. In our experiment, isolation does not ensure self-fertilization as snails were old enough to have had breeding encounters prior to isolation. To determine the impact of antibiotics on the rate of development of the parasite and infection prevalence in snails, parasite production (count of parasite larvae [cercariae]) was assessed weekly in *B. glabrata* beginning week 4 post exposure until the end of the experiment at week 9. This window coincides with a 4-week pre-patent period of parasite development where no parasite release occurs, followed by release of cercariae beginning at approximately week 4.

To measure parasite production, snails were placed in well plates with 10mL of well water and positioned under fluorescent light to allow parasite emergence. After 1 hour, snails were returned to their respective jars and the presence or absence of cercariae was recorded. If cercariae were detected, a 1 mL aliquot of well water was taken and all cercariae within the aliquot counted (Gleichsner et al., 2016). Finally, the survival of snails was checked weekly.

### Statistical analysis

Data on parasite and host reproduction had considerable zero-inflation. As such, we constructed mixed effects hurdle models to account for data overdispersion using the glmmTMB package in R (Brooks et al. 2017). Hurdle models first model the probability of obtaining a zero-value, similar to logistic regression. Then, if the value is non-zero, hurdle models use a specific error distribution to further predict host/parasite reproduction. As such, all hurdle model outputs contain coefficients for a zero-inflated model, predicting the probability of a zero, and a conditional model, predicting non-zero measurements based on a specific error distribution. Models were fit with treatment, week of experiment, and treatment * week of experiment interactions terms as fixed effects except where inclusion of the interaction term was uninformative. Host individual was used as a random intercept within each model to account for repeated measures on individual host snails. However, no random slope (week of experiment) was incorporated as individual snails within a treatment show similar temporal patterns in host and parasite development at this time scale. Additionally, we generated these hurdle models using both a Poisson and negative binomial error distribution for non-zero values and used AIC to determine which model best fit the data. Statistical models were visualized using the effects package in R 3.6.3.

To examine parasite development time, we conducted a time-to-event analysis (survival analysis) to determine how long post exposure infected snails would begin producing parasites. Additionally, a survival analysis was run on all individuals within a treatment, irrespective of infection status, to look for differences in survival among treatments. Examining all individuals within each treatment was necessary as absence of infection cannot be definitively confirmed until week 6, meaning infected individuals who died prior to shedding would be unknown. Both survival analyses were run in R 3.6.3 using the survival and survminer packages. Analysis was conducted over the course of the 9-week experiment and used to parameterize daily mortality rates within our mathematical.

### Mathematical model

In order to further understand how antibiotic contamination may alter disease dynamics, we adapted the differential equation model of Hoover et al. (2020). We used a base model without predation or agrochemical pollution, including snail reproduction associated with fecundity compensation (Minchella and LoVerde, 1981; Figures S1-S3) and delayed castration, and assumed all female *S. mansoni* worms are paired. These alterations result in the set of differential equations and dynamic variables below. Here, S, E, and I represent susceptible, exposed, and infected snails, respectively. W represents the mean worm burden in the human population with M representing the number of female worms, C representing infective parasite cercariae, and N the total snail population:

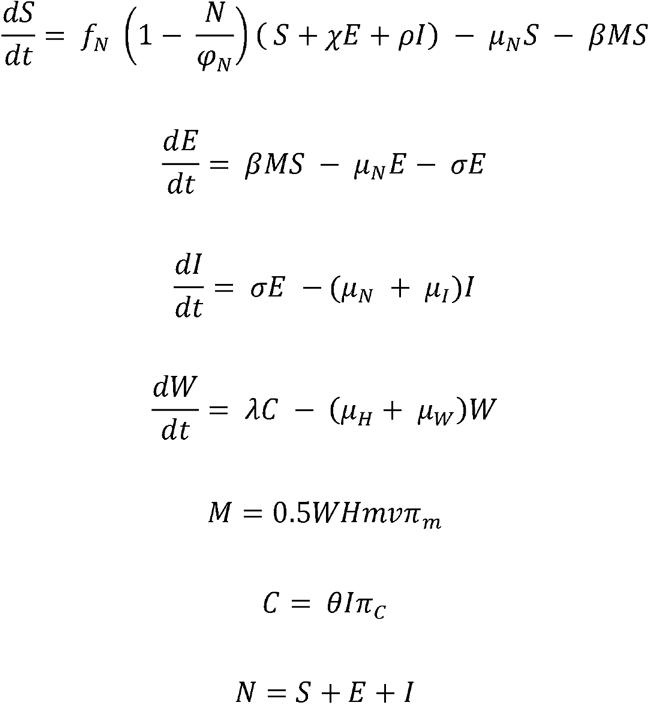

We used our experimental data to calculate the percent change in snail and parasite life history characteristics in response to antibiotic exposure and altered model parameters accordingly (Table S2). This exercise is intended only to make qualitative predictions on how antibiotic exposure may alter disease dynamics. Analysis was run using the deSolve package in R 3.6.3 (Soetaert et al., 2010).

## Results

### Parasite Production

Antibiotics accelerated parasite development. Antibiotic + parasite snails released parasites on average one week earlier than parasite only snails (Survival analysis χ^2^= 8.9, p = 0.003, Figure 1). Although significantly earlier parasite release occurred in the antibiotic + parasite treatment, the number of cercariae released was low. As such, the impact of early maturation may be limited. Infection prevalence for the antibiotic + parasite and parasite only treatments were not significantly different at 77% and 59%, respectively (two-sample equality of proportion test, χ^2^= 1.696, p = 0.193). Additionally, antibiotics had a dynamic effect on parasite production over time such that the antibiotic + parasite treatment had lower initial parasite production but produced more parasites on average over the course of the experiment. (Figure 2 and Table 1, see Figure S4 in the supplement for raw data visualization).

**Figure 1.**
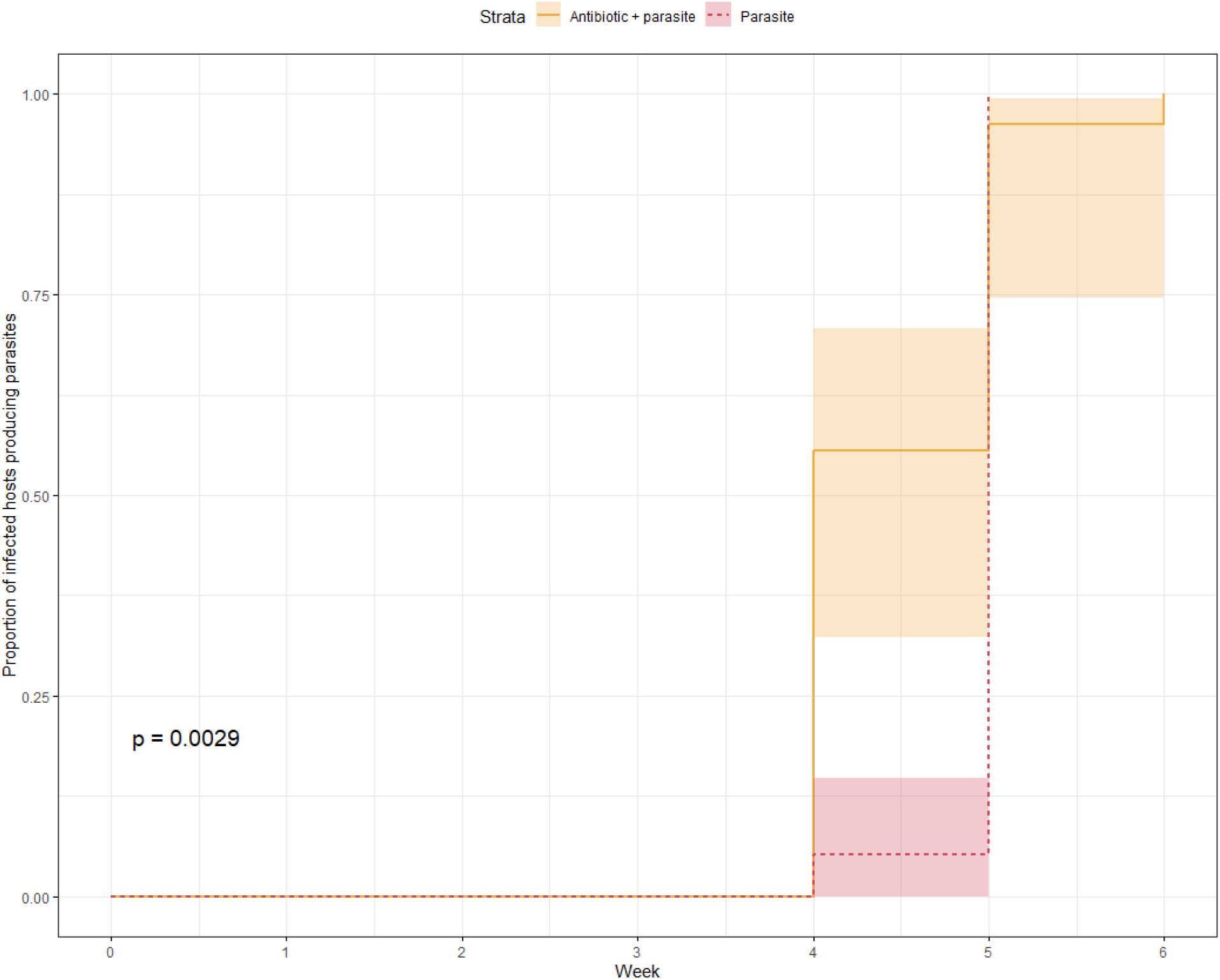
Time from initial exposure until snail hosts began producing parasites. The proportion of *Schistosoma mansoni*-infected snails that released parasite cercariae in the parasite only treatment (red) and the antibiotic + parasite treatment (gold) was significantly different (χ^2^= 8.9, p = 0.0029) with antibiotic + parasite snails releasing parasites earlier than parasite only snails.

**Figure 2.**
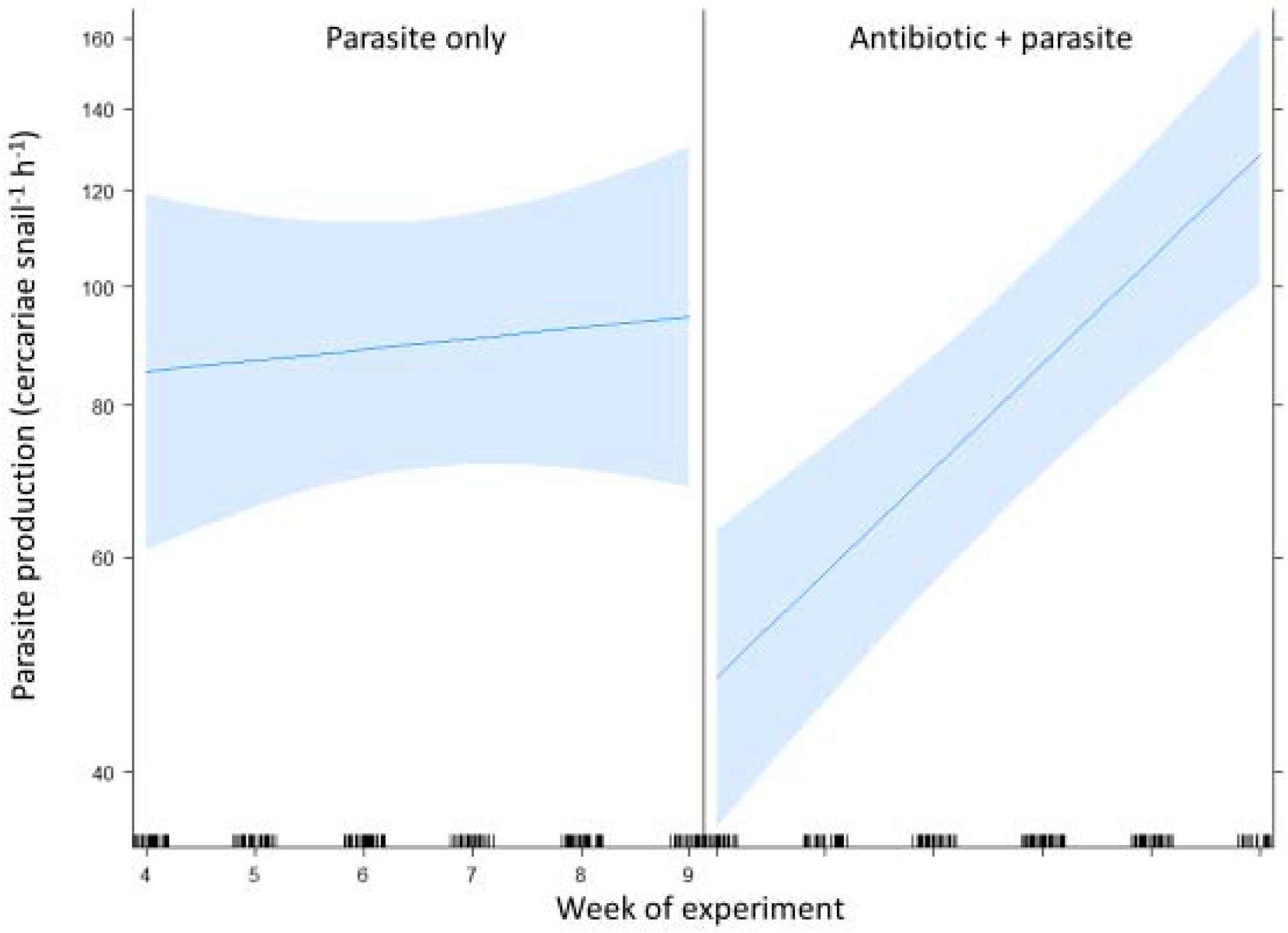
Parasite production (cercariae per snail per hour) during the patent period of infection in the experiment (week 4 – week 9). Antibiotic + parasite snails had initially low parasite production that increased as the infection progressed compared to the parasite only treatment. The y axis is on a log scale to account for negative binomially distributed data. See Table 1 for summary statistics.

**Table 1.** Model results for parasite production comparing the antibiotic + parasite treatment relative to the parasite treatment (intercept). Coefficients of the zero-inflation model show the probability of obtaining a zero value.

| **Zero Inflation Model** | **Coefficient** | **Standard error** | **Z value** | **P value** |
| --- | --- | --- | --- | --- |
| Intercept | 3.841 | 1.180 | 3.255 | 0.001 * |
| Antibiotic + parasite | 1.765 | 2.267 | 0.778 | 0.436 |
| Week | -0.867 | 0.218 | -3.982 | <0.001 * |
| Antibiotic + parasite: Week | -0.519 | 0.471 | -1.102 | 0.270 |
| **Conditional Model** |  |  |  |  |
| Intercept | 4.362 | 0.334 | 13.069 | < 0.001 * |
| Antibiotic + parasite | -1.285 | 0.417 | -3.083 | 0.002 * |
| Week | 0.021 | 0.047 | 0.441 | 0.660 |
| Antibiotic + parasite: Week | 0.177 | 0.058 | 3.058 | 0.002 * |

### Host Reproduction

Snails in the antibiotic treatment were more likely to lay eggs relative to snails in the control treatment (Zero-inflation component of Table 2, p = 0.005, Figure 4, Figure S5). The antibiotic + parasite treatment had a higher probability of laying eggs than the parasite only treatment throughout the entire experiment (Zero-inflation component of Table 3, p = 0.006). Additionally, comparison of the parasite treatment with the antibiotic + parasite treatment suggests an initially similar reproductive output (Figure 3), yet as the infection matured, snails within the antibiotic + parasite treatment had a higher probability of laying eggs. These snails evinced delayed castration compared to the parasite only treatment snails (visible as wide confidence intervals for the parasite treatment in Figure 3, also see the zero-inflation component of Table 3). The observed decrease in eggs laid over time from infected snails in both treatments is due to parasitic castration (Minchella and LoVerde, 1981).

**Table 2.** Model results for host reproduction comparing the antibiotic treatment relative to the control treatment (intercept). Coefficients of the zero-inflation model show the probability of obtaining a zero value.

| **Zero Inflation Model** | **Coefficient** | **Standard Error** | **Z Value** | **P Value** |
| --- | --- | --- | --- | --- |
| Intercept | -0.812 | 0.540 | -1.503 | 0.132 |
| Antibiotic | -2.166 | 0.700 | -2.813 | 0.005 * |
| Week | -0.121 | 0.0587 | -2.062 | 0.039 * |
| **Conditional Model** |  |  |  |  |
| Intercept | 1.107 | 0.095 | 11.676 | <0.001 * |
| Antibiotic | 0.190 | 0.105 | 1.813 | 0.070 |
| Week | 0.030 | 0.010 | 2.890 | 0.004 * |

**Table 3.** Model results for host reproduction comparing the antibiotic + parasite treatment relative to the parasite treatment (intercept). Coefficients of the zero-inflation model show the probability of obtaining a zero value.

| **Zero Inflation Model** | **Coefficient** | **Standard Error** | **Z Value** | **P Value** |
| --- | --- | --- | --- | --- |
| Intercept | -7.756 | 1.688 | -4.596 | <0.001 * |
| Antibiotic + parasite | 4.667 | 1.702 | 2.742 | 0.006 * |
| Week | 2.609 | 0.523 | 4.989 | <0.001 * |
| Antibiotic + parasite:<br>Week | -1.820 | 0.505 | -3.603 | <0.001 * |
| **Conditional Model** |  |  |  |  |
| Intercept | 1.705 | 0.220 | 7.733 | <0.001 * |
| Antibiotic + parasite | 0.337 | 0.252 | 1.338 | 0.181 |
| Week | -0.201 | 0.111 | -1.812 | 0.070 |
| Antibiotic + parasite:<br>Week | -0.061 | 0.121 | -0.502 | 0.616 |

**Figure 3.**
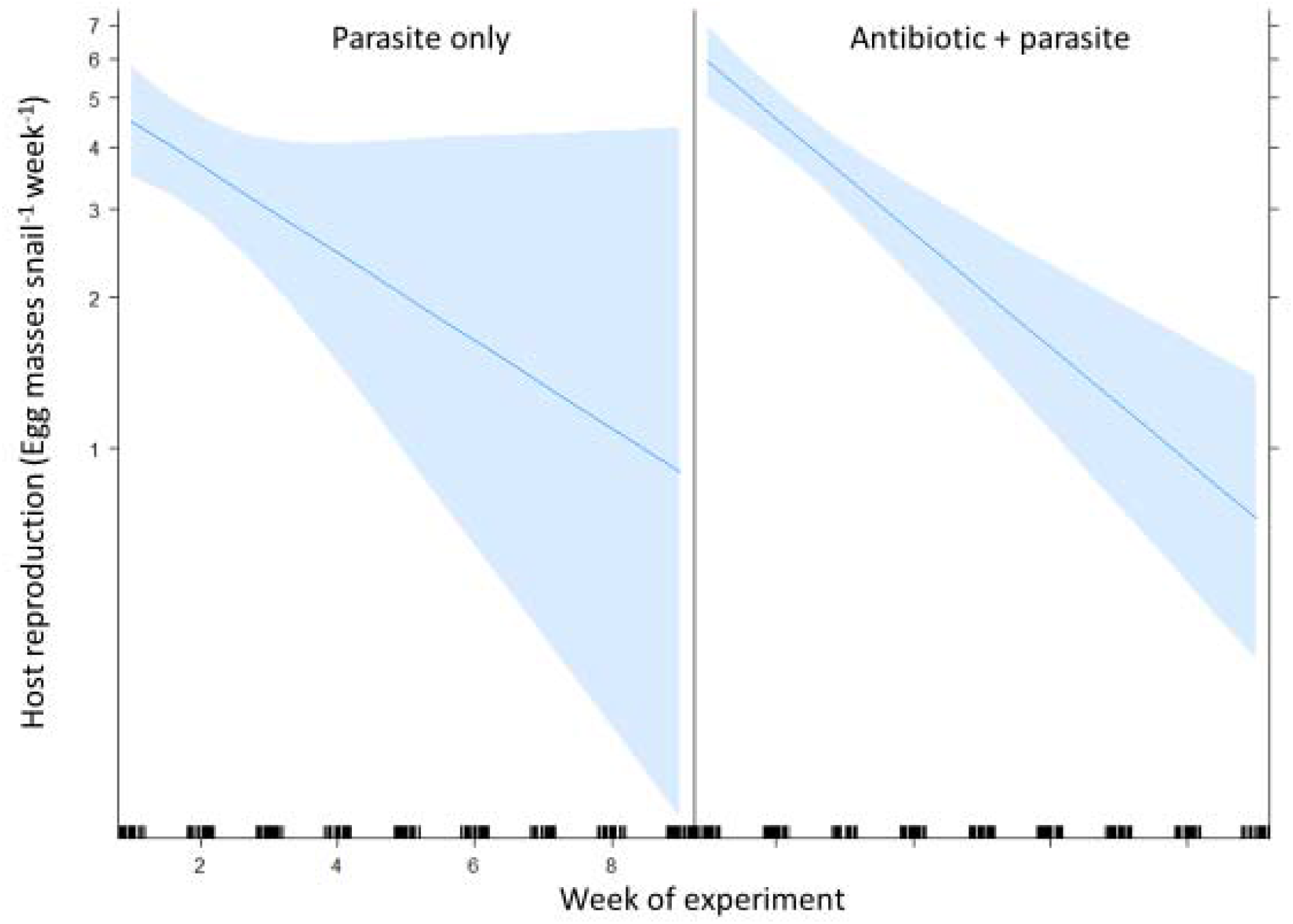
Host reproduction (egg masses per snail per week) over the course of the experiment (week 1 – week 9). Antibiotic + parasite and parasite only treatments have initially similar host reproductive patterns. However, after week 4, parasite only snails were more likely to lay no eggs than antibiotic + parasite snails as evinced by wide confidence intervals in the parasite only treatment. The y axis is on a log scale to account for negative binomially distributed data. See Table 3 for summary statistics.

**Figure 4.**
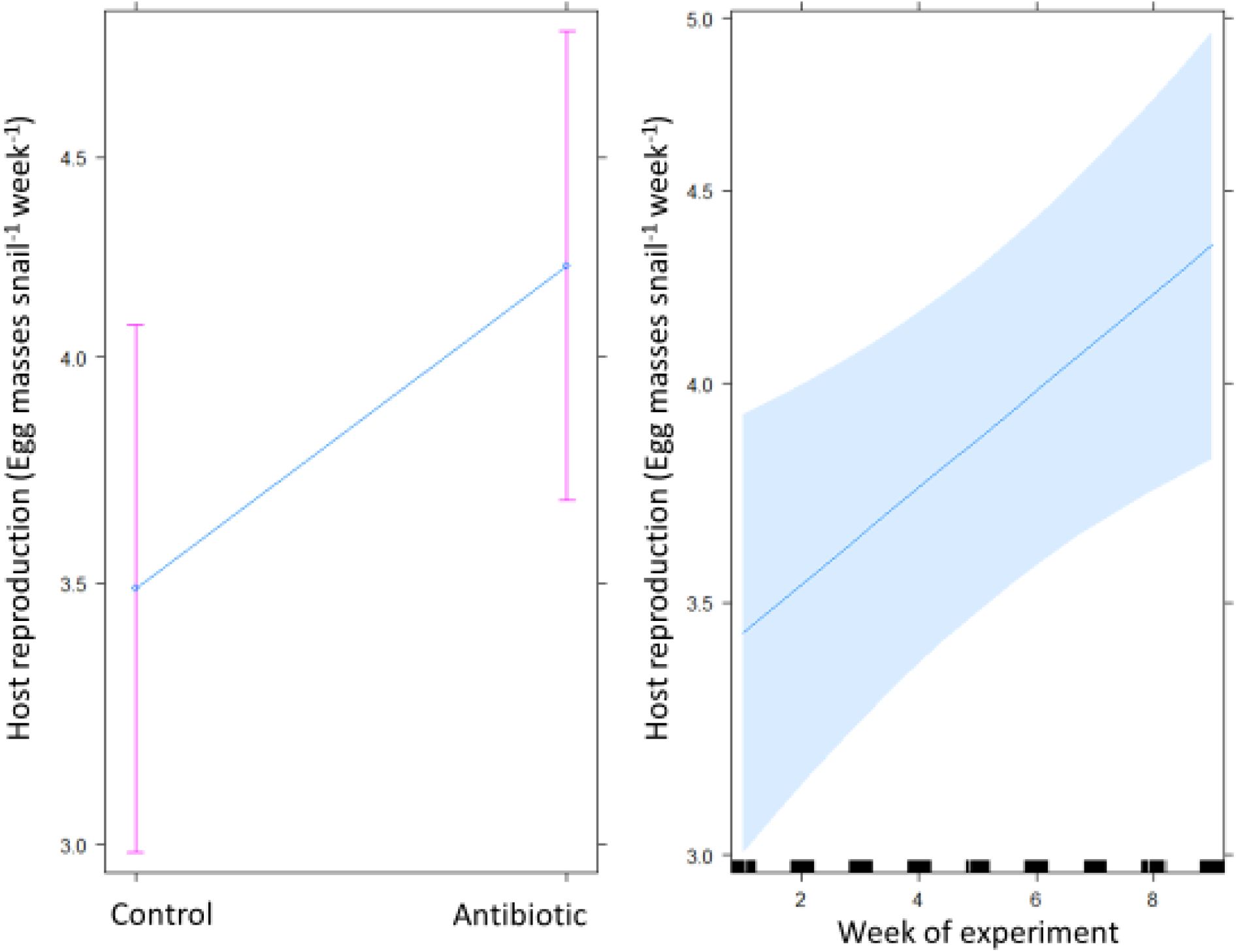
Host reproduction (egg masses per snail per week) over the course of the experiment (week 1 – week 9). Antibiotic only snails were significantly more likely to produce offspring compared to control snails. Additionally, snails were more likely to produce offspring and produced more offspring later in the experiment. The interactive effect of treatment * week was uninformative in this statistical model and was omitted. See Table 2 for summary statistics.

The antibiotic only treatment showed the highest survival, followed sequentially by the control treatment, then parasite only treatment, and antibiotic + parasite treatment (see supplemental Figure S6). However, only the antibiotic + parasite versus the parasite only survival curves were significantly different from one another (χ^2^=5.4, p = 0.02).

Based on our adaptation of a published *S. mansoni* model, our model results suggest that areas with antibiotic contamination may have increased snail exposure to parasites and more rapid snail population growth, leading to higher infection prevalence in snails and greater worm burdens in humans (Figure 5). Additionally, snail exposure and human worm burdens increase more rapidly in the antibiotic scenario than would otherwise occur (see supplemental Figure S7 for proportional changes in model state variables).

**Figure 5.**
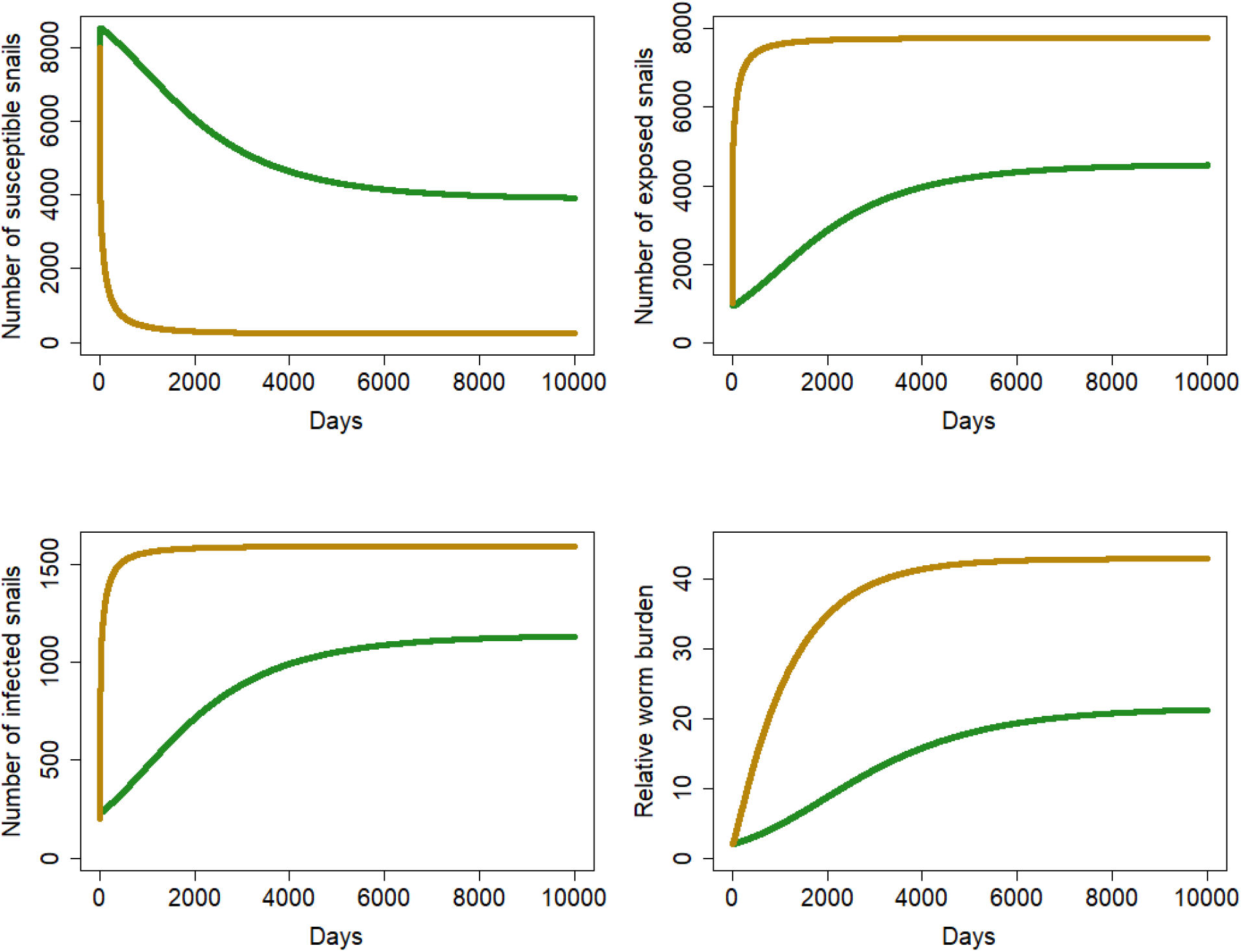
Model output showing the number of susceptible snails, exposed snails, infected snails, and mean worm burden in the human population in the antibiotic (gold; data from antibiotic only and antibiotic + parasite snails) versus control (green; data from control and parasite only snails) scenarios. Antibiotics are likely to increase *Schistosoma* infections in snails and humans based on parameterizations generated from our experimental data.

## Discussion

We investigated the impact of an ecologically relevant concentration of tetracycline on the life history parameters of the trematode parasite *S. mansoni* and its snail intermediate host, *B. glabrata.* We assessed host reproduction and survival as well as parasite production and development time-in the presence and absence of the antibiotic, tetracycline. Our results suggest that antibiotics are likely to impact snail and parasite production with potentially significant ecological ramifications. We show that tetracycline facilitated earlier parasite production by infected hosts and increased parasite output as the infection matured (Figures 1 and 2, respectively). Additionally, the presence of antibiotics increased egg laying in uninfected snails when compared to uninfected, well water controls (Table 2). Lastly, parasitic castration is delayed in the antibiotic + parasite snails, and these snails had a significantly higher egg output throughout the experiment compared to the parasite only treatment (Table 3). To the best of our knowledge, this is the first study to document the impact of antibiotic contamination on the host and parasite life history parameters of this freshwater snail and its medically relevant parasite.

Modifications in host-parasite interactions by antibiotic contamination are likely associated with changing microbiome dynamics (Hernández-Gómez et al., 2020, Knutie et al., 2017) and correspond to our hypothesized mechanism. Antibiotics often disturb microbiomes by decreasing useful and/or increasing harmful bacteria (Akbar et al., 2020). Of the four most common bacterial genera found in the flora of 200 snails, *Acinetobacter* and *Vibrio* are both responsive to tetracycline, suggesting a biological linkage between antibiotic presence and infection susceptibility is possible in this system (Portet et al., 2021). The alterations observed in parasite development time, parasite production, and host production from addition of tetracycline are congruent with research findings of microbiome-related impaired immune function following parasitic infection (Portet et al., 2018). Portet et al. (2018) proposed that following infection with *S. mansoni*, the bacterial microbiome of *B. glabrata* changed its composition, which could account for the altered immune function. Decreased immune function weakens defense mechanisms within hosts (Akbar et al., 2020), which can increase susceptibility to parasites and corresponds with our findings of an accelerated parasite development time in the antibiotic + parasite treatment (Pravdová et al., 2020). A potential alternative explanation is that the parasites were more aggressively exploiting their host, indicating increased parasite pathogenicity (Anzia and Rabajante, 2018, Schlüter-Vorberg and Coors, 2019). Our results suggest that aggressive exploitation may have been occurring as snails showed significantly lower survival in the antibiotic + parasite treatment compared to the antibiotic control. However, distinguishing between these two mechanisms will require further study.

Another finding consistent with an altered microbiome was the significant increase of parasite production in the antibiotic + parasite treatment compared to the parasite only treatment in the final weeks of the experiment. A study conducted by Schlüter-Vorberg and Coors (2019) found similar results when studying the effects of pharmaceuticals on pathogen virulence. Schlüter-Vorberg and Coors (2019) demonstrated that chemically induced suppression of immune systems in *Daphnia* weakens disease resistance by enhancing the virulence of the parasite and increases the proportion of infected hosts. However, they additionally observed an increased speed of host sterilization, which contrasts with our findings of a delayed castration period in hosts. Our initial survival data suggest that the observed increase in reproductive output may also be associated with increased mortality within infected snails in the antibiotic + parasite treatment, which could result in a net decrease in parasite fitness. However, our model suggests that the increased mortality in infected snails is not sufficient to balance the other life history alterations associated with antibiotic exposure. Considering all factors, parasite transmission and thus disease burden in humans are likely to increase in regions with sufficient antibiotic concentrations.

Finally, hosts exposed to antibiotics were more likely to lay eggs. Increased reproduction due to antibiotic exposure has also been shown in other studies of invertebrates (Flaherty and Dodson, 2005). A similar study investigating the effect of pharmaceuticals on *Daphnia* reproduction found that chronic exposure to certain types of antibiotics induced significantly faster development and more reproduction (Flaherty and Dodson, 2005). These influences on typical development patterns varied depending on both the duration of exposure, and number of pharmaceuticals to which they were exposed. The strongest and most complex effects between fecundity and antibiotics were observed with long term (30-day) exposure and a combination of multiple pharmaceuticals mixtures, respectively. In contrast, inhibition of growth and reproduction have been recorded using a 100 μg/mL solution of the antibiotic streptomycin (Chernin, 1957). These contradicting results may have arisen from the drastic 2000-fold difference in antibiotic concentration used. Our results, which emphasize the potential of antibiotics to alter host-parasite population dynamics, may have wider implications for food webs as snails are important herbivores and prey within many systems (Johnson, 2009). The capacity of antibiotics to change food web dynamics could also manifest as an increased antibiotic sensitivity in primary producers such as algae, affirming the potential risks of contamination in non-target species that could additionally influence snail-host populations (Isidori et al., 2005). For example, tetracyclines are known to impact ecosystems via cytotoxic effects that limit plant growth and reduce chlorophyll content (Polianciuc et al., 2020). The findings presented here support the idea that ecologically relevant tetracycline concentrations accelerate parasite development time, increase reproductive output in parasites, and enhance host reproduction, possibly through antibiotic-induced changes to the microbiome.

## Conclusion

We are only beginning to understand the impacts of antibiotics on hosts, parasites, and their interactions. Here, we show that antibiotics influence parasite dynamics by facilitating earlier parasite production with increasing output as the infection matures. Infected hosts affected by antibiotic contamination demonstrated increased egg laying and egg output throughout the experiment when compared to the parasite only treatment. In addition, parasite castration was delayed in hosts exposed to antibiotics. Our study suggests that the continued widespread use of antibiotics with improper disposal results in residual consequences to freshwater ecosystems and may increase the burden of schistosomiasis in endemic regions. The largely unknown ecological and anthropogenic impacts of antibiotic contaminants including - but not limited to - trophic effects, disease risk, and ecosystem interactions therefore merit further research. As antibiotic usage increases, its role as a link between human health and host-parasite interactions emphasizes the need to further explore the consequences of human activity on all facets of global change.

## Acknowledgements

We would like to thank members of the Minchella lab for providing feedback and assistance in preparing this manuscript. This research did not receive any specific grant from funding agencies in the public, commercial, or not-for-profit sectors.

